# Transcriptional Profiling Reveals the Transcription Factor Networks Regulating the Survival of Striatal Neurons

**DOI:** 10.1101/2020.05.20.105585

**Authors:** Lin Yang, Zihao Su, Ziwu Wang, Zhenmeiyu Li, Zicong Shang, Heng Du, Guoping Liu, Dashi Qi, Zhengang Yang, Zhejun Xu, Zhuangzhi Zhang

## Abstract

The striatum is structurally highly diverse, and its organ functionality critically depends on normal embryonic development. Although several studies have been conducted on the gene functional changes that occur during striatal development, a system-wide analysis of the underlying molecular changes is lacking. Here, we present a comprehensive transcriptome profile that allowed us to explore the trajectory of striatal development and identify the correlation between the striatal development and Huntington’s disease (HD). Furthermore, we applied an integrative transcriptomic profiling approach based on machine learning to systematically map a global landscape of 277 transcription factor (TF) networks. Most of these TF networks are linked to biological processes, and some unannotated genes provide information about the corresponding mechanisms. For example, we found that the Meis2 and Six3 were crucial for the survival of striatal neurons, which was verified using conditional knockout (CKO) mice. Finally, we used RNA-Seq to speculate their downstream targets.

## Introduction

The striatum is the major information processing centre in the basal ganglia circuits of the mammalian forebrain. It is involved in multiple neurological functions ranging from motor control, cognition, emotion, and reinforcement to plasticity underlying learning and memory ^1^. The dysfunction of neurotransmission or neurodegeneration of the striatum results in a number of neurological disorders, including Huntington’s disease (HD), Parkinson’s disease (PD) and obsessive–compulsive disorders ^2–4^. Thus, exploring the striatal development may provide new insights for the treatment of these diseases.

Transcription factors (TFs) and genetic regulatory networks (GRNs) are essential for the differentiation, migration and survival of striatal neurons. Currently, most studies have focused on a few genes through specific individual pathway to regulate striatal development. Based on these efforts, the key roles of several transcriptional factors (Gsx2, Dlx1/2, Sp8/9, Isl1, etc.) in controlling striatal development have been explored ^5–7^. Although our knowledge of striatal development has been enhanced by substantial data from reports, their limited focus hampers our understanding of the whole process in the context of systems biology. In fact, TFs are highly dynamic and interactive, and TF networks regulate striatal development through highly coordinated expression changes ^8^. Thus, mapping the global landscape of TF networks during striatal development may provide some insights at the system level. Genes that belong to the similarity functional network may be co-expressed in the temporal logic and, thus, should have similar expression profiles to a certain extent ^9–11^. Correlation analysis between these expression profiles have become the premise of the guilt-by-association approach, which is used to identify genes co-expressed with known genes (probes) to achieve a specific function or expression pattern ^12^. This strategy of using a priori knowledge and data integration can further enhance the elucidation of gene regulatory functions ^12^. By using known functional annotations as labels for given genes, one can determine the extent to which the network summarizes the information by using a machine learning model to classify these genes as belonging to a provided function based on their correlation in the co-expression network ^13^.

In the present work, we analysed the developmental process of the striatum from a macro-scale perspective and found that HD was closely related to the later stage of striatal development. Furthermore, we provided the global landscape of 277 TF networks in striatal development based on machine learning, and these TF networks can be used to infer potential transcription factor functions. For example, our results showed that Six3 and Meis2 were included in the TF networks involved in neuronal apoptosis processes. Using conditional knockout (CKO) mice, we verified that Six3 and Meis2 are indeed involved in the neuron apoptotic processes during striatal development. These results further validate the rationality of our global landscape of TF networks. Furthermore, we predicted the downstream targets of Six3 and Meis2 by analysing the RNA-Seq data of the Six3 and Meis2 CKO mice, respectively. In conclusion, this study provides global TF networks, which offer valuable assets and provide direct insight into the molecular details of the striatal developmental process, enabling us to better understand the striatal development and its related disease.

## Results

### Systematic Analysis of Striatal Development Based on Transcriptomes

To investigate the changes of gene expression during striatal development, we performed RNA-Seq from the early stage to the adult stage (Fig. 1). After calculating the normalized gene expression levels as the fragments per kilobase of the transcript per million mapped reads (FPKM; See Methods), we first performed a correlation analysis of the experimental data between 5 time points. We found that all data showed good consistency and a high degree of correlation (R = 0.95–0.99) between the temporally adjacent experiments (Fig. S1A). After pre-processing the raw data (See Methods), a list of 15443 genes was compiled from all 5 time points (Fig. S1B; Table S1).

**Fig. 1.**
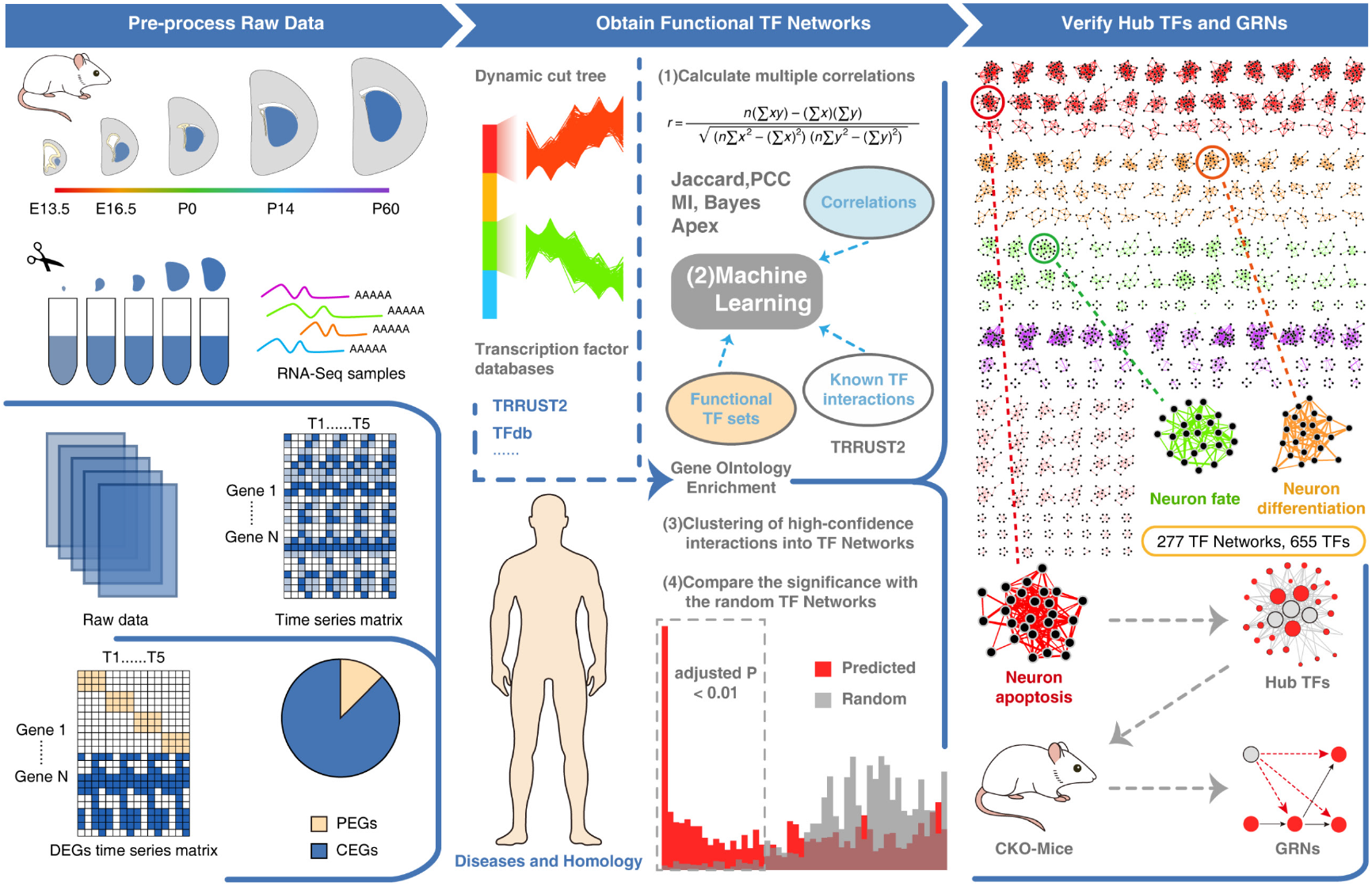
Overview of the experimental design. Firstly, 5 time-points of striatum mRNA measured during mouse development using RNA-Seq. Then, the raw transcriptome data were pre-processed to obtain the DEGs. The DEGs were divided into two parts (See Methods): PEGs and CEGs. Secondarily, the functional TF networks of CEGs were predicted based on co-expression analysis and machine learning. And then the homologous and disease genes were annotated from existing databases. Finally, the functional significance of these TF networks was tested. Next, the hub TFs in the related TF networks were predicted based on the ARACNE algorithm. And then, CKO mice were constructed to verify the hub TFs and infer their GRN.

To further investigate the developmental status of the striatum from E13.5 to P60, we obtained the differentially expressed genes (DEGs) (See Methods). We identified all 8861 DEGs, including 882 (10%) TFs (Fig. 2A; Table S2), and then performed biological process and KEGG pathway enrichment analysis to assign the genes to functional categories (Table S2). The functions of these DEGs were significantly enriched in biological processes, including the “regulation of cell cycle” (E13.5-E16.5), “regulation of synapse organization and axonogenesis” (E16.5-P0), “regulation of ion transmembrane transport” (P0-P14), “positive regulation of intrinsic apoptotic signalling pathway and ATP synthesis” (P14-P60) and other biological processes (Fig. S2A). The functions of these DEGs were significantly enriched in pathways including the “cell cycle and Hippo signalling pathway” (E13.5-E16.5), “axon guidance and cAMP signalling pathway” (E16.5-P0), “calcium signalling pathway and MAPK signalling pathway” (P0-P14), “Huntington disease and Alzheimer disease” (P14-P60) and other pathways (Fig. S2B). Throughout the whole process of striatal development, our results indicated that cell division gradually decreased. In contrast, neuronal differentiation, synapse organization and the dopamine signalling pathway were gradually enhanced from E13.5 to P14 (Fig. 2B). Interestingly, we found that the significantly altered genes from P14 to P60 are strongly associated with Huntington’s disease in the striatum (Fig. 2C). Therefore, our transcriptome dataset and a collected transcriptome dataset of HD (See Methods) obtained from 38 patients^14^ enabled us to analyse the correlation between striatal development and HD at the mRNA level. To correlate striatal development with HD at the level of pathways, we listed all gene products that were detected in two studies and counted the number of significantly altered gene products in one of the seven pathways known to be involved in development and HD (Fig. 2D; Table S3). Two-tailed hypergeometric statistical tests (See Methods) were carried out to determine whether there was significant overlapping of altered gene products on a specific pathway between striatal development and HD (Fig. 2E and 2F). Importantly, all pathways exhibited high correlations (−log_10_P-value > 1.3). Hence, these analyses provided evidence that striatal development was closely associated with HD processes.

**Fig. 2.**
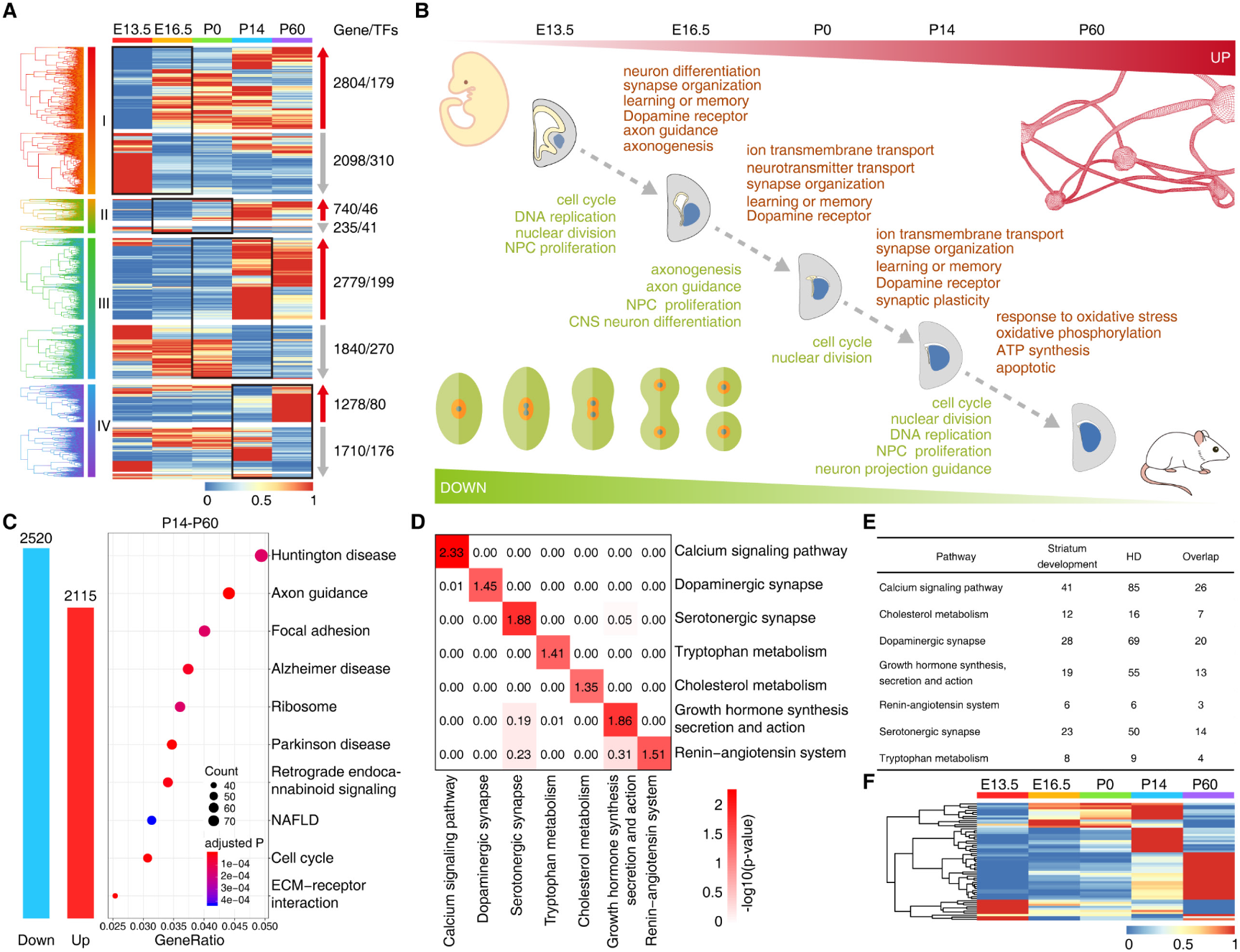
Striatal development and correlation analysis with HD. **A** Expression patterns of DEGs between two adjacent time points. The number of genes and TFs are shown on the right. **B** The patterning of striatal development in mice. **C** Histogram of DEGs (FDR < 0.05) in the striatum from P14 to P60. The top 10 significant KEGG pathways of these DEGs are displayed. **D** Two-tailed hypergeometric tests were conducted to determine whether there is a significant overlap of altered genes on a specific pathway between striatal development and HD. Correlations on seven pathways between striatal development and HD. **E** The number of significantly altered and overlapping genes on seven signaling pathways in striatal development and HD. The genes (FDR < 0.05) differentially expressed in striatal development and HD (adjusted P < 0.05) **F** Temporal expression patterns of 57 overlapping genes across 5 time-points.

### Co-expression Analysis Identified Interrelated Functional Modules of Continuously Expressed Genes

To further provide insights into the functional transitions during striatal development, we divided the sum of DEGs obtained into periodically expressed genes (PEGs) and continuously expressed genes (CEGs) (See Methods; Table S4). We used all 1257 PEGs, including 115 (9%) TFs from E13.5 to P60, and then performed biological process enrichment to assign the genes to functional categories (Fig. S3A and S3B). Our results show that the PEG biological process is in line with our delineative striatal development model in Fig. 2B. We applied weighted gene co-expression network analysis (WGCNA) ^15^ to obtain a system-wide understanding of groups of CEGs (7095 genes, including 740 TFs) (Table S5) whose co-expression patterns are highly correlated during striatal development. The soft-threshold power of WGCNA was defined as 13 and the scale-free topology index was > 0.85 (Fig. S4A). In total, 6 distinct co-expression modules were identified that showed a diverse trend of gene expression, and each module was assigned a colour name (Fig. 3A; Table S5). Then, to show the trends in gene expression, the genes from each module were grouped into two trend clusters using the fuzzy c-means algorithm ^16^ (Fig. 3A; See Methods). Clearly, each module contained not only positively correlated genes but also negatively correlated genes. Next, the network heatmap plot further suggested those main modules were independent from each other (Fig. S4B).

**Fig. 3.**
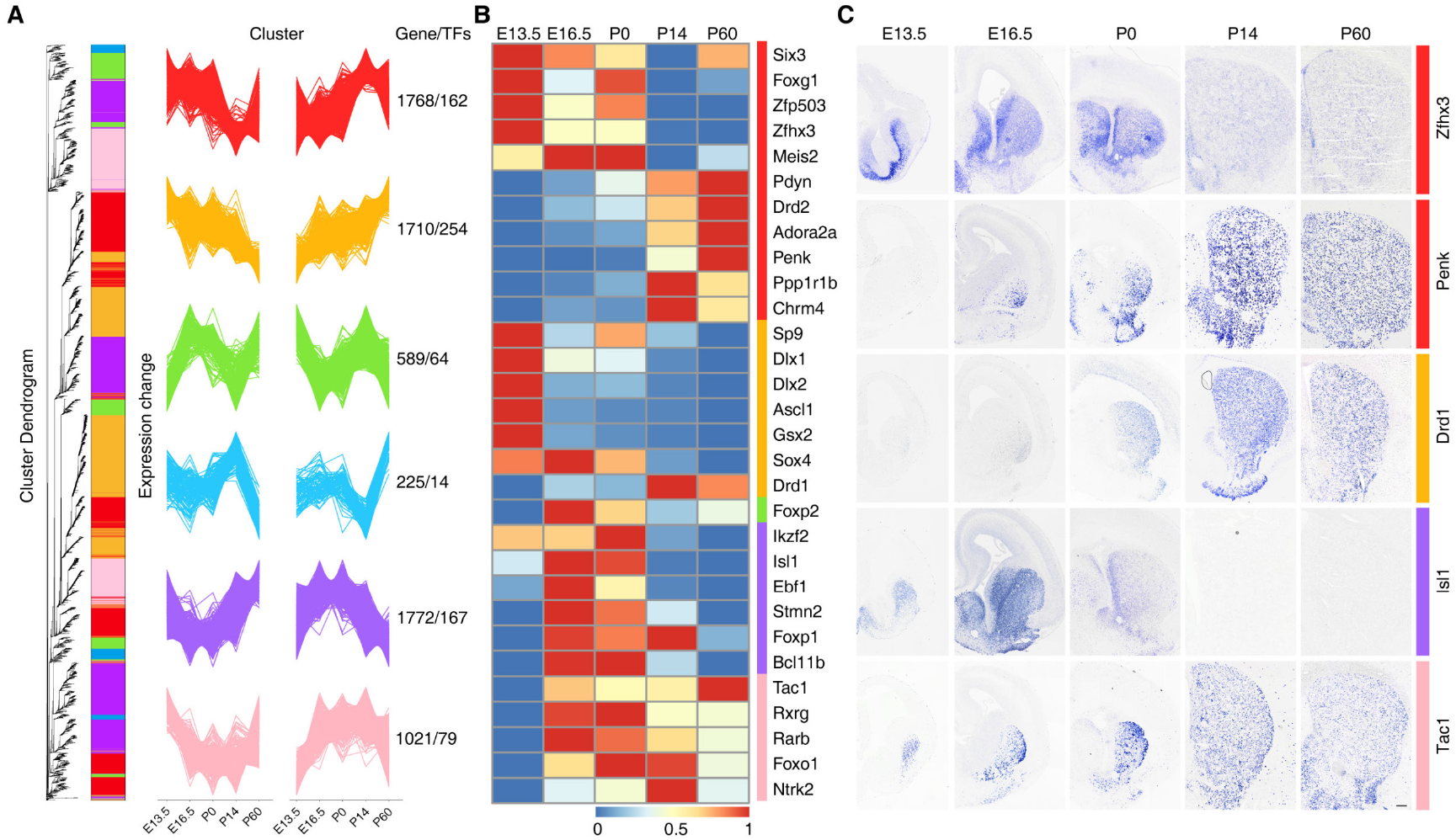
Co-expression analysis of differentially expressed continuous genes. **A**Hierarchical clustering dendrogram based on the dynamic hybrid tree cut algorithm shows co-expression modules of continuous genes that are colour coded. The co-expressed genes are rearranged, and fuzzy c-means clustering identifies 2 distinct temporal patterns of each co-expression module. The x axis represents five developmental points, while the y axis represents log2-transformed, normalized intensity ratios. The number of genes and TFs in each module are shown on the right. **B** Expression pattern of reported striatal development genes at 5 time-points. **C** The expression patterns of striatal development genes were verified by in situ hybridization at 5 time-points. Scale bars: 100 μm in c.

In addition, biological process and KEGG enrichment analysis of the gene members in each module revealed distinct biological functions and pathways (Fig. S5A and S5B; Table S5). Intriguingly, genes in the orange module were continuously increased or decreased at the 5 time points (Fig. 3A). Genes in this module are enriched in “mitotic nuclear division” and “DNA replication” (Fig. S5A and S5B), which is consistent with cell division and neuronal differentiation during striatal development. Many genes are essential for striatal MSN proliferation and differentiation, including Gsx2, Dlx1/2 and Sp9, which were defined in this module (Fig. 3B). Genes in the red module were enriched in pathways related to “synapse organization”, “regulation of transmembrane transporter activity”, and “neurotransmitter transport”, and all were related to synaptic transmission (Fig. S5A). Some genes were essential for striatal development, including Zfhx3, Adora2a, Ppp1r1b and Drd2, which were defined in this module (Fig. 3B). Finally, we selected genes randomly and verified the expression trends of these genes (Zfhx3, Penk, Drd1, Tac1 and Isl1) by in situ hybridization (Fig. 3C). Our results showed that the expression trends of these genes were correct according to our co-expression modules. Altogether, these results not only make it possible to understand the dynamic events during striatal development but may also help to identify key factors.

### Global Map of Potential Functional TF Networks Based on Machine Learning

From our previous analysis, we identified transcriptional regulators that exhibited clear temporal dynamics. At the core of the processes that regulate striatal development and function is the transcriptional circuitry. This prompted us to focus on TF expression in our time series. We identified 740 differentially expressed TFs from six CEG co-expression modules (Fig. 4A; Table S5), and these TFs were used to predict functional TF networks based on machine learning. Using the co-expression of TFs with optimized parameter settings and known functional sets (Table S6), machine learning was performed by the following steps (See Methods): correlation metrics were calculated (five algorithms), machine learning predicted interactions, and high-confidence interactions were clustered into functional networks (Fig. S6A). The output of machine learning was well organised and ready for use in subsequent analysis to predict functional TF networks (Fig. 4B). The distribution of the interaction network node degree distribution followed a power-law (Fig. S6B), which has certain biological significance, and this phenomenon is also widely confirmed ^17^. The high proportions of human homologous TFs ^18^ and nervous system disease TFs ^19^ in the mice involved in these networks suggested that the subsequent analysis was likely to provide a reference for human striatum studies (Fig. S6C and S6D). Afterward, we parted the network using the ClusterONE algorithm to partition the TF networks for downstream analysis ^20^. To assess the physiological significance of the putative striatum functional networks, we analysed the TF networks for coherent biological functions based on GO annotations. We found that almost all of the TF networks in striatal development were significantly enriched for associations with essential processes (Table S7). To prove that the predicted TF networks obtained through machine learning have more biological significance than just co-expression analysis, we used the method of comparison with a random model to assess the accuracy of the predicted TF networks (See Methods). For each module, which was measured as the biological process enrichment adjusted p-value distribution, there was markedly higher significance for the function of putative TF networks compared to the same number of randomized networks of TFs in the network that were not predicted (Fig. 4C and S7). Therefore, 277 putative TF networks were identified from five co-expression modules (Fig. 4D; Table S7).

**Fig. 4.**
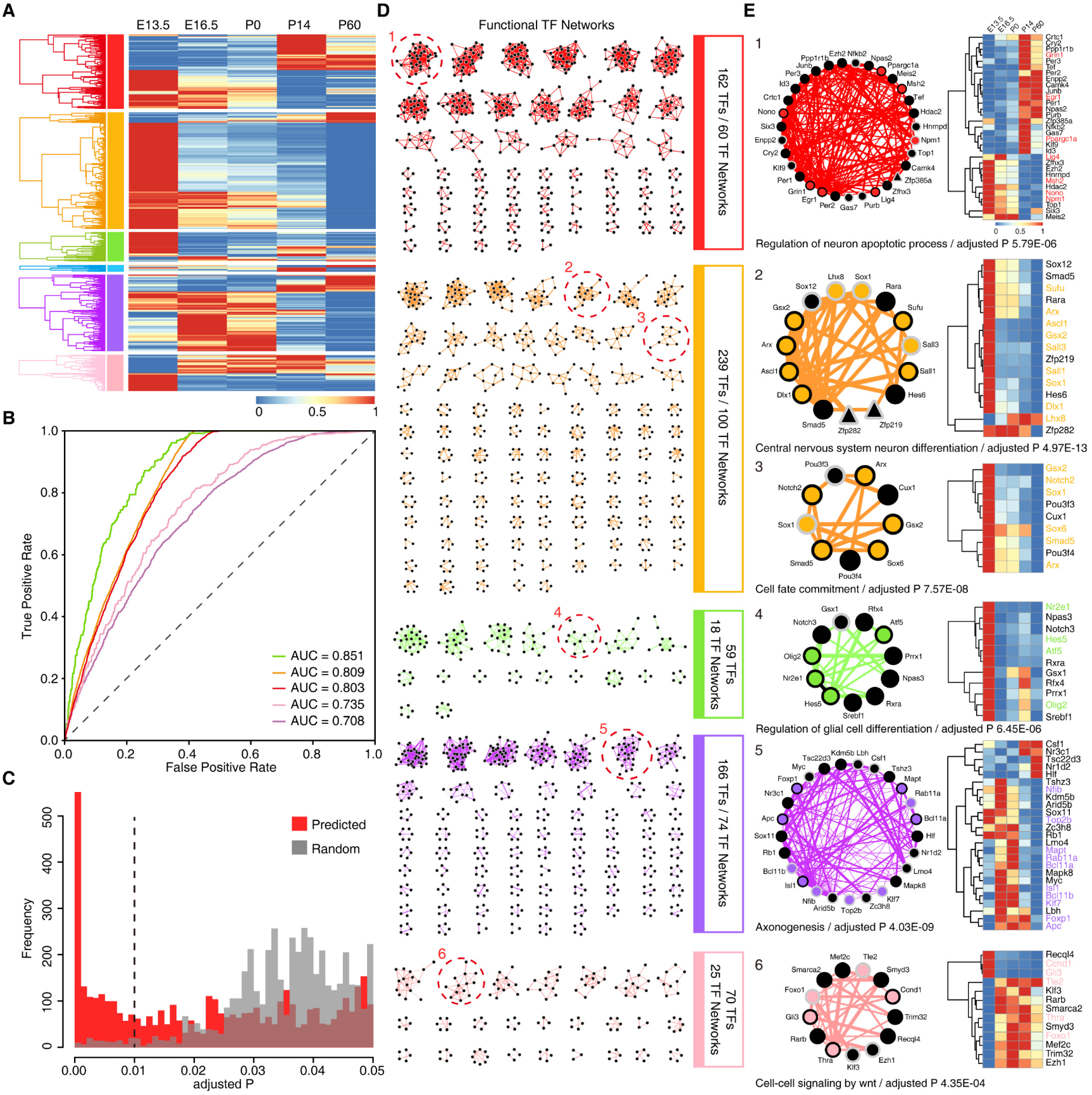
Predicted potential functional TF networks based on machine learning. **A** Expression patterns of 740 differentially expressed continuous TFs from different co-expression modules. **B** Receiver operating characteristic of differentially expressed continuous TFs from each co-expression modules. The AUC is used to assess the quality of machine learning. The AUC of the diagonal is 0.5. **C** Functional significance distribution of predicted TF networks and random TF networks. **D** Schematic of the global atlas of inferred functional TF networks in striatal development. **E** Example of TF networks with different functions of known and unknown subunits. The width of the edges increases with increasing correlation score. The shape of the node represents whether it is a homologous TF (circular node for homologous to humans, triangle node for unique in mouse). The colour of the node border represents whether the TFs are associated with nervous system–associated disease (black border for disease TFs, grey border for non-disease TFs). The black node indicates potential functional TFs. Expression patterns of representative TF networks are shown on the right by heatmap.

Afterward, we roughly predicted some novel TF functions. For example, we predicted that TF network 1 was enriched in the “neural apoptosis process” with high significance (adjusted P 9.85E-06) (Fig. 4E). Likewise, we predicted that TF network 2 was enriched in “central nervous system neuron differentiation” with high significance (adjusted P 4.97E-13) (Fig. 4E). In this TF network, these TFs (Ascl1, Gsx2, Dlx1, Lhx8, Sox1, Sufu, Arx, Sall1 and Sall3) have been reported to be associated with cell differentiation ^5,21–29^. We also predicted other functional TF networks enriched in “cell fate commitment”, “axonogenesis”, and “cell-cell signalling by wnt” (Fig. 4E). These functional TF networks can provide us with rich treasures for predicting novel functional genes, which will be helpful for researchers carrying out work related to the striatum, including exploring the core transcriptional regulators and constructing transcriptional regulatory networks.

### CKO of putative TFs leads to abnormal apoptosis in striatal neurons

Neuronal apoptosis is very important in the development of the nervous system, and the striatum is no exception. Abnormal neuronal apoptosis leads to many neurological diseases, such as HD and Parkinson's disease, which are closely related to the striatum. Therefore, it is necessary to explore neuronal apoptosis during striatal development, which will provide new insights and ideas for researchers. Next, we mainly verified the predicted neuronal apoptosis-related TFs to further confirm that our global landscape of TF networks was reliable. The TF network mentioned above (Fig. 4E), is significantly enriched for biological processes including three main categories “neuron apoptotic process” (Fig. 5A) based on the GO term relationships using BiNGO^30^. To test this hypothesis, we selected several core factors: Zfhx3, Six3 and Meis2 (Fig. 5B), in the center of the network, all of which were poorly studied in “regulation of neuron apoptotic process” in the striatum. Recently, the TF Zfhx3 was reported to induce abnormal cell apoptosis in the striatum during the first postnatal week ^31^. However, whether Six3 and Meis2 are involved in the cell apoptosis of striatal MSNs is largely unknown.

**Fig. 5.**
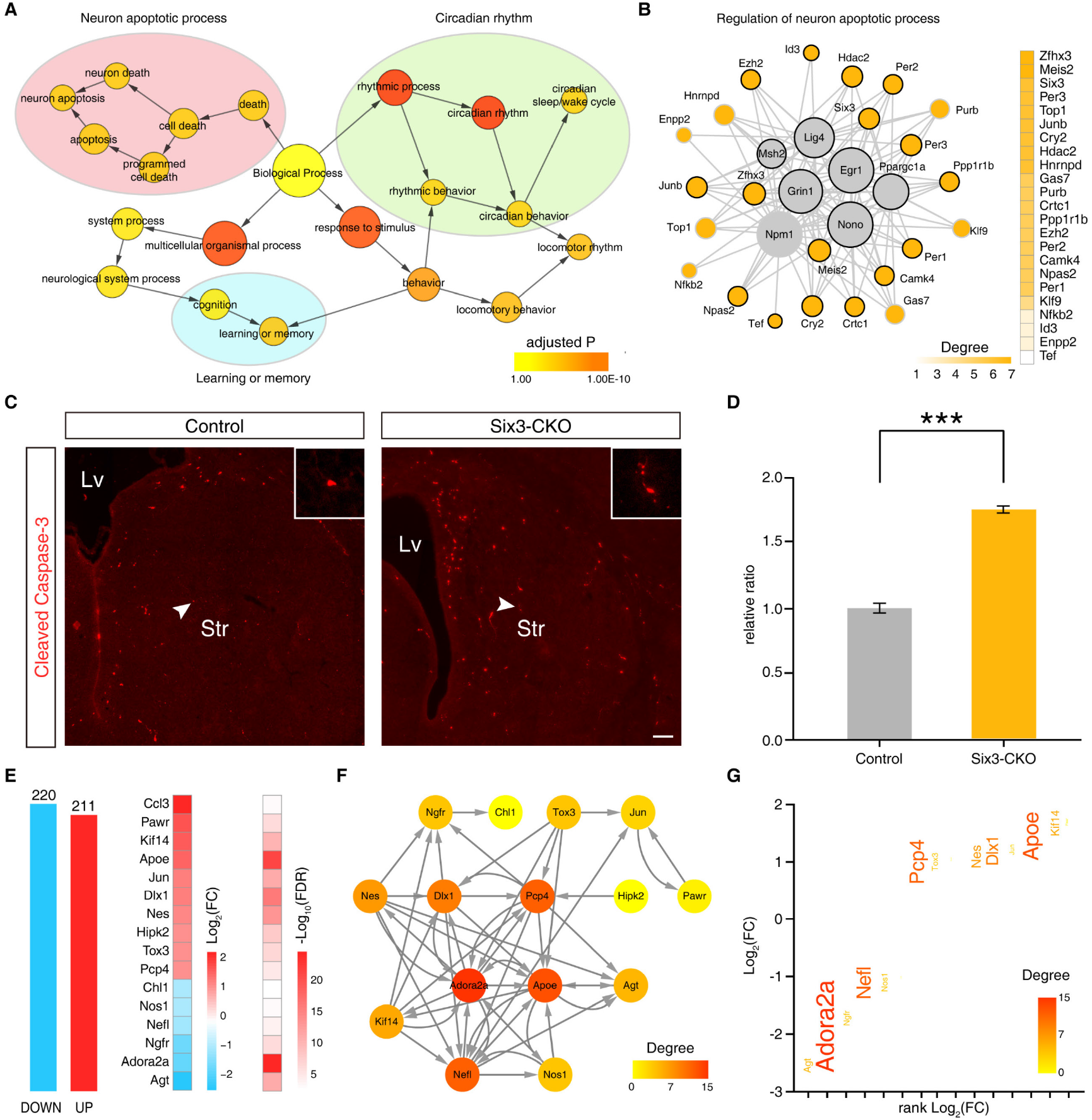
Novel functions of Six3 predicted from TF networks. **A** Part of the BiNGO results for a TF network, as visualized in Cytoscape. Red categories are the most significantly overrepresented. Yellow nodes are not significantly overrepresented, which include orange nodes in the context of the GO hierarchy. The area of a node is proportional to the number of genes in the test network annotated to the corresponding GO category. **B** Topological representation of the reconstructed “regulation of neuron apoptotic process” TF regulatory network using ARACNE. TFs are represented as nodes and inferred interactions as edges. Grey nodes indicate the known functional TFs. Potential functional TFs are coloured orange. The degree of orange nodes is shown on the right. **C** A significant increase in expression of cleaved Caspase-3 in Six3-CKO striatum compared to controls at P0. Scale bars: 100 μm in c. **D** Histogram showing that the number of cleaved Caspase-3^+^ cells was increased in Six3-CKO mice. (Student’s t-test, ***P < 0.001, n=3 mice per group, mean ± SEM). Lv: lateral ventricle. Str: striatum. **E** Histogram of DEGs in the LGE at P0 between control and Six3-CKO mice. The 16 DEGs associated with “regulation of neuron apoptotic process” are shown. **F** A GRN of these “regulation of neuron apoptotic process” genes was inferred by Genie3. The edge direction represents the regulatory interaction from regulatory genes to target genes. Node size and colour are proportional to the number of genes (in-degree and out-degree). **G** Key downstream genes that are candidates for regulating the neuronal apoptotic process. The significantly changed genes in Six3-CKO mice versus wild-type mice are shown. Genes are coloured by their importance (degree) in the “regulation of neuron apoptotic process” GRN, and the font size increases with the degree of node.

To investigate whether Six3 plays a functional role in striatal MSN apoptosis, we conditionally deleted the Six3 gene in the LGE using Dlx5/6-Cre lines (henceforth described as Six3-CKO). Cre recombinase was robustly expressed in the LGE and Six3 was removed from the striatum ^32^. Next, we performed immunostaining of cleaved Caspase-3 at P0. Our results showed that the cleaved Caspase-3 positive cells were increased approximately 2-fold in the Six3-CKO mice compared with their littermate controls (Fig. 5C, D). To investigate the potential mechanism by which Six3 affects neuronal apoptosis in the striatum, we performed RNA-Seq analysis at P0 between Six3-CKO mice and their littermate controls. Our RNA-Seq data revealed that approximately 431 DEGs in the Six3-CKO mice (220 downregulated and 211 upregulated RNA expression) and then performed biological process enrichment of these DEGs to functional categories (Fig. 5E, Supplementary Table 8), suggesting that Six3 plays a key role in regulating neuronal apoptosis of the LGE at P0 (regulation of neuron apoptotic process/ adjusted P: 5.45E-05/ 16 DEGs). To further explore the potential hub downstream factors for regulating neuronal apoptosis of Six3, we constructed the GRN of 16 significantly changed genes using Genie3 (see Supplementary Methods), which predicted the directed regulatory links from genes to genes (Fig. 5F). To determine which genes might be controlled by Six3 in the striatum, we also measured the static network parameter of this GRN. Adora2a was thought to have high pleiotropic effects and play essential roles in regulating neuronal apoptosis (Fig. 5G). Changes in Adora2a were expected to have larger biological impacts than those occurring in genes in the network periphery. This suggests that Adora2a may be a potential hub downstream factor for Six3 to regulate neuronal apoptosis. Adora2a is highly expressed in D2-type MSNs and is involved in cell apoptosis^33,34^. These findings suggest that Six3 regulated striatal MSN apoptosis by regulating the expression of Adora2a.

To further verify the rationality of our global landscape of TF networks and confirm whether Meis2 was involved in striatal MSN apoptosis, we designed the Meis2 targeting vector to generate the Meis2^Flox^ allele through the strategy of homologous recombination (Fig. 6A). To analyze whether Meis2 plays a functional role in MSN apoptosis, we used Dlx5/6-Cre to conditionally knockout Meis2 (henceforth described as Meis2-CKO) in the striatum. Our results showed that Meis2 mRNA expression was completely lost in the LGE or striatum in Meis2-CKO mice compared with that in their controls by in situ hybridization (Fig. 6B). Note that the expression of Meis2 in the striatum was completely lost but the expression of Meis2 in the cortex remained. Next, we performed immunostaining of cleaved Caspase-3 at P0. There were more cleaved Caspase-3 positive cells in the Meis2-CKO mice than in their controls (Fig. 6C, D). In order to investigate the potential mechanism of Meis2 affecting neuronal apoptosis in the LGE or striatum, we performed RNA-Seq analysis at E16.5 between Meis2-CKO mice and their littermate controls. Our RNA-Seq revealed that approximately 858 DEGs in Meis2-CKO mice (632 downregulated and 226 upregulated RNA expression) and then performed biological process enrichment of these DEGs to functional categories (Fig. 6E; Supplementary Table 9), suggesting that Meis2 plays a key role in neuronal apoptosis of the striatum (regulation of neuron apoptotic process/ adjusted P: 9.65E-05/ 22 DEGs). To further explore the potential hub downstream factors for regulating neuronal apoptosis of Meis2, we constructed the GRN of 22 significantly changed genes using Genie3 (see Supplementary Methods), which predicted the directed regulatory links from genes to genes (Fig. 6F). To determine which genes might be controlled by Meis2 in the striatum, we also measured the static network parameter of this GRN. Isl1 and Adora2a were thought to have high pleiotropic effects and play essential roles in regulating neuronal apoptosis (Fig. 6G). Changes in Isl1 or Adora2a were expected to have larger biological impacts than those occurring in genes in the network periphery. Our results suggest that Isl1 and Adora2a may be potential downstream factors for Meis2 to regulate the neuronal apoptosis. In fact, Isl1 and Adora2a have been reported to be expressed in D1-type and D2-type MSNs and regulate cell apoptosis in the striatum^6,33,34^. Altogether, these results indicated that the transcriptional factors Six3 and Meis2 are the hub genes in the transcriptional network for the survival of striatal MSNs.

**Fig. 6.**
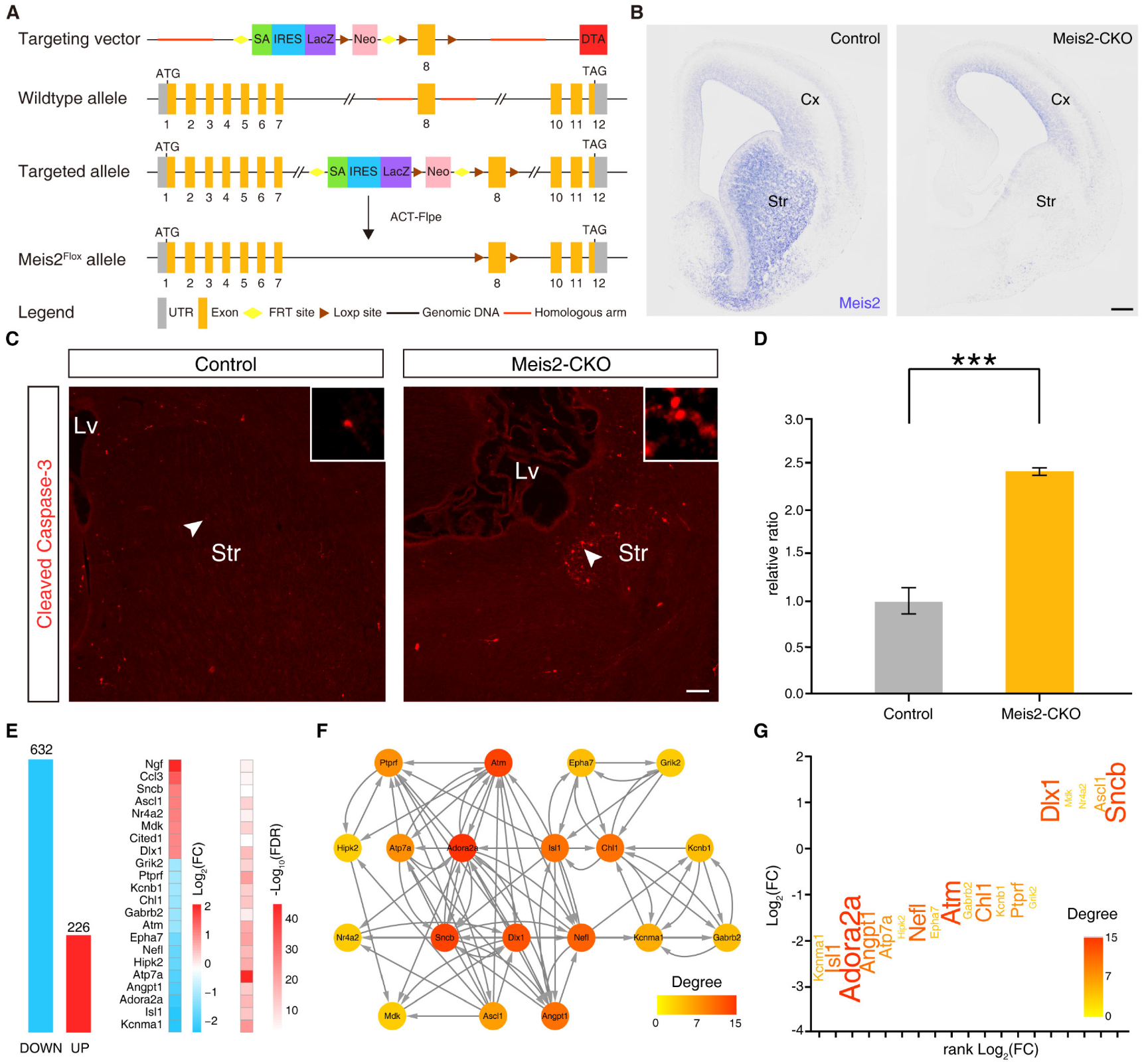
Meis2 regulates neuronal apoptotic processes in the striatum. **A** The generation of the Meis2^Flox^ allele. The Meis2 gene-targeting vector was designed to delete exon 8 of Meis2. **B** Deletion of the Meis2 gene was confirmed at the mRNA level by in situ hybridization. **C** The immunofluorescence staining of cleaved Caspase-3 in Meis2-CKO striatum and controls at P0. Scale bars: 100 μm in b and c. **D** Histogram showing that cleaved Caspase-3^+^ cells were increased in Meis2-CKO at P0. (Student’s t-test, ***P < 0.001, n = 3 mice per group, mean ± SEM). Cx: cerebral cortex. Str: striatum. **E** Histogram of DEGs in the LGE between control and Meis2-CKO mice at P0. The 18 DEGs associated with “regulation of neuron apoptotic process” are shown. **F** A GRN of these “regulation of neuron apoptotic process” genes was inferred by Genie3. The edge direction represents the regulatory interaction from regulatory genes to target genes. Node size and colour are proportional to the number of genes (in-degree and out-degree). **G** The significantly changed genes in Meis2-CKO mice compared to control mice are shown which are candidates for regulating the neuronal apoptotic process. Genes are coloured by their importance (degree) in the “regulation of neuron apoptotic process” GRN, and the font size increases with the degree of node.

## Discussion

In this study, we presented the panoramic overview of the developing mouse striatum from embryonic stage to adult. Quantitative analysis of 15443 gene products allows us to explore the trajectory of striatal development and identify the correlation between the striatal development and HD. More importantly, we provided the global landscape of 277 TF networks based on co-expression analysis and machine learning models. Furthermore, we identified the hub TFs Six3 and Meis2 which are involved in regulating the apoptosis of striatal neurons. Finally, using conditional knockout (CKO) mice and RNA-Seq data, we verified that Six3 and Meis2 indeed regulated neuronal apoptosis and inferred their downstream targets in the striatal development stages. Therefore, our study provides a resource to understand striatal development and its connection to striatal disease.

During mouse striatal development, our preliminary bioinformatic analysis suggested that cell division gradually decreased, but neuronal differentiation, synapse organization and the dopamine signalling pathway gradually increased. It is worth mentioning that the DEGs in the striatum from P14 to P60 are closely related to HD. This phenomenon inspired us to investigate whether there is experimental evidence for a connection between striatal development and HD. By comparing the mRNA expression of striatal development with that of HD ^14^, we found that a noticeable number of pathway regulators in HD are also differentially expressed during striatal development. Moreover, we identified statistically significant networks of overlapping genes in seven pathways between striatal development and HD, which suggests that HD regains some features of striatum embryonic development but loses some basic functions of the adult striatum and that HD is likely a consequence of the dysregulation of master regulators and pathways.

The main motivation behind this study was the identification of networks of co-expressed TFs, with an inherent expression distribution corresponding to discrete biological processes associated with striatal development. Firstly, individual pair-wise TF associations were scored based on the temporal expression profile similarity measured during striatal development. Next, we used an integrative computational scoring procedure to derive TF interactions. The support vector machine learning classifier used was trained on correlation scores obtained for reference annotated TF sets ^35^, and we combined all of the input co-expression data with previously published TF interaction evidence ^36^. Measurements of the overall performance showed a high area under the curve for the TF co-expression data. The final filtered 277 TF networks consists of 655 transcription factors. From these TF networks, we identified transcriptional regulators across all networks, including Ascl1 Gsx2 and Dlx1, which were highly co-expressed in the LGE during the development of the striatum, along with their co-expressed targets. Deletion of the expression of these genes leads to abnormal differentiation of striatal MSNs ^5,22,28^. Thus, these TF networks can be used to speculate novel gene functions during striatal development. For example, we identified two key factors (Six3 and Meis2) from one TF network related to neuronal apoptosis. Since neuronal apoptosis is one of the major events in striatal development, we further explored the possible downstream factors of these novel functional genes to elucidate their potential mechanisms underlying neuronal apoptosis. We generated Meis2-CKO mice for the first time. Using Meis2-CKO mice to perform RNA-Seq, we predicted that Isl1 and Adora2a were regulated by Meis2 in regulating the neuronal apoptosis. These findings were supported by previous reports ^6,33,34^. Moreover, Using the same approach, we speculated that Six3 regulated the survival of striatal neurons by regulating the expression of Adora2a. Adora2a is highly expressed in D2-type MSNs and is involved in cell apoptosis ^33,34^. This raises the exciting potential to identify more critical factors from our data and deepen our understanding of the regulation of striatal development.

## Methods

### Animals and Tissue Collection

The mouse genetic background was CD1 in this study. The Dlx5/6-Cre and Six3^F/F^ mice were previously described ^21,37^. We generated Meis2^F/+^ mice in this study. The details of the targeting construct are shown in Fig. 6. Briefly, we first generated Meis2 knockout mice via the ESC targeting system. The cassette is composed of an FRT site followed by a LacZ sequence and a loxP site. This first loxP site is followed by neomycin under the control of the human beta-actin promoter, SV40 polyA, a second FRT site and a second loxP site. A third loxP site is inserted downstream of the targeted exon 8. Exon 8 is thus flanked by the loxP site. A “conditional ready” (floxed) allele can be created by flp (ACT-flp) recombinase expression in mice carrying this allele. Subsequent Cre expression results in knockout mice. The Mesi2^F/F^ mouse genotype was identified by PCR (Polymerase Chain Reaction). The primers were as follows: Meis2-loxp F1: TGTCAAACATTGCCATCCAAACAGT, and Meis2-loxp F2: TCATGGAGAAGACAGCGCTGGT. Mice were group-housed with approximately three mice in each cage on a 12-h light, 12-h dark cycle, with ad libitum chow and water. The day of vaginal plug detection was calculated as E0.5, and the day of birth was considered as P0. Both males and females were used in all experiments. All animal experiments described in this study were approved in accordance with institutional guidelines at Fudan University Shanghai Medical College.

RNA-Seq analysis was performed as previously described ^38^. For the temporal transcriptome analysis, the whole LGE or striatum was separated from embryonic to adult mice. The LGE or striatum was collected at 5 time points: E13.5, E16.5, P0, P14 and P60. Each time point had three biological replicates, but we obtained only two successful biological replicates at P14. The whole LGE or striatum tissues were washed twice with ice-cold HBSS (Hanks’ Balanced Salt Solution) to remove blood, quick-frozen in liquid nitrogen and then stored at −80 °C for total RNA extraction. The striatum (including the VZ, SVZ and MZ-striatum) of the Dlx5/6-Cre; Six3^F/F^ (henceforth described as Six3-CKO) mice and their Dlx5/6-Cre littermates (henceforth described as controls) were dissected at P0 (n=3 mice, each group). The LGE or striatum (including the VZ, SVZ and MZ-striatum) of the Dlx5/6-Cre; Meis2^F/F^ (henceforth described as Meis2-CKO) mice and their Dlx5/6-Cre littermates (henceforth described as controls) were dissected at E16.5 (n=3 mice, each group).

### Immunohistochemistry

Mice were anaesthetized with an intraperitoneal injection of ketamine (100 mg per kg) and transcardially perfused with a fixative solution containing 4% paraformaldehyde (PFA). The brains were post-fixed in 4% PFA at 4 °C overnight. Sections of 20 μm thickness were used for immunostaining. The sections were washed with 0.05 M TBS for 10 min, incubated in Triton-X-100 (0.5% in 0.05 M TBS) for 30 min at room temperature (RT), and then incubated with block solution (10% donkey serum +0.5% Triton-X-100 in 0.05 M TBS, pH=7.2) for 2 h at RT. Primary antibodies against cleaved Caspase-3 (1:500; Cell Signaling, 9661, 43) were diluted in 10% donkey serum block solution, incubated overnight at 4 °C, and then rinsed 3 times with 0.05 M TBS. Secondary antibodies (from Jackson, 1:500) matching the appropriate species were incubated for 3 h at RT. The fluorescently stained sections were then washed 3 times with 0.05 M TBS. This was followed by 4′, 6-diamidino-2-phenylindole (DAPI) (Sigma, 200 ng/ml) staining for 3 min, and the sections were then cover-slipped with Gel/Mount (Biomeda, Foster City, CA). Fluorescence images were collected with an Olympus-BX61VS microscope using 10× and 20× objectives.

### In Situ RNA Hybridization

In situ RNA hybridization experiments were performed using digoxigenin labelled riboprobes on 20-μm frozen sections as previously described ^31,32^ or made from cDNAs amplified by PCR using the following primers:

Drd1 Fwd: GGCATTTGGAGAGATGTGGCACCAG.
Drd1 Rev: AGATAGCCCAATACCTGTCCACGCT.
Meis2 Fwd: CGATGGGTTAGACAACAGCGTAGCTTCACCTG.
Meis2 Rev: ATGGCTTGGCAAATATGATGCATTGGGTCCATG.
Isl1 Fwd: ATGGGAGACATGGGCGATCC.
Isl1 Rev: CATGCCTCAATAGGACTGGCTACC.
Penk Fwd: TCGGAAGGACAGGATGTCATCA.
Penk Rev: CGTCAGGAGAGATGAGGTAACAAAC.
Tac1 Fwd: CCCCTGAACGCACTATCTATTC.
Tac1 Rev: TAGAGTCAAATACCGAAGTCTCAG
Zfhx3 Fwd: CGATCTGGCCCAGCTCTACCA.
Zfhx3 Rev: CTGTAAGCCTGCGAGGGCATAG.

### Quantification

DAPI staining was used to define the striatal area for quantification. Brain regions were identified using a mouse brain atlas and sections equivalent to the following coordinates: the most-rostral section, 2.19 mm; the most-caudal section, 3.03 mm, n=3 mice for each genotype, 4 sections per mouse). Schema chart was shown in Figure S8. For quantification of Caspase-3 positive cells in entire striatum at P0, four 20-μm thick coronal sections from rostral, intermediate, and caudal levels of the striatum were analysed. We counted all Caspase-3 positive cells in the striatum.

### RNA Sequencing and Data Processing

The total RNA was extracted from the LGE or striatum tissues and treated with deoxyribonuclease I (DNase I). mRNA was isolated by Oligo Magnetic Beads and cut into small fragments that served as templates for cDNA synthesis. Once short cDNA fragments were purified, they were extended with single-nucleotide adenines, ligated with suitable adapters, and amplified by PCR before they were sequenced. High-throughput RNA sequencing (RNA-Seq) experiments were carried out using the Illumina comprehensive next-generation sequencing (NGS) technique. The raw data were filtered and processed by FastQC software (Version 0.11.5, available online at the website: http://www.bioinformatics.babraham.ac.uk/projects/fastqc). Low-quality RNA-Seq reads were removed if they had a Phred quality score of less than 20 or had less than 50 nucleotides. The filtered reads were mapped onto the mouse reference genome (GRCm38.p2.genome, released on 12/10/2013) using HISAT2 software (Version 2.1.0). Assembly and quantification of the transcripts were accomplished with StringTie software (Version 1.3.1) using the mouse genome annotation file as the reference (gencode.vM2. annotation, available at the GENCODE website). The number of FPKM was used for the measurement of the relative abundances of the transcripts.

The Pearson correlation coefficient was calculated between biological replicates with the normalized expression levels of log_2_ (FPKM value +1). The missing value was defaulted to 0. The comparison of the two biological replicates showed that the expression values between them were highly correlated (average R = 0.98). Hence, the average FPKM value of the two replicates was taken as the expression level for the sample at each time point. However, prior to this point, the transcription was performed at least twice in biological repetitions. Afterward, to reduce the influence of transcription noise, a gene was defined as expressed if its FPKM value was ≥1 at least at a time point.

The read counts were utilized for differential expression analyses using the edgeR package in R. A key feature of the edgeR package is the use of weighted likelihood methods to implement a flexible empirical Bayes approach in the absence of easily tractable sampling distributions ^39^. Gene expression data between adjacent time points (E13.5-16.5, E16.5-P0, P0-P14 and P14-P60) were used for differential analysis. The false discovery rate (FDR) was controlled by the Benjamini and Hochberg (BH) algorithm. Unless otherwise specified, genes with a FDR < 0.01 and a fold-change (FC) > 2 or < 0.5 were judged to be differentially expressed. Also, the DEGs were divided into two parts: PEGs (the genes differentially expressed between only two adjacent time points) and CEGs (the genes differentially expressed between at least three adjacent time points).

Hierarchical clustering was performed by the pheatmap package in R. Additionally, the transformed and normalized gene expression values were calculated by dividing their expression levels at different time points by their maximum observed FPKM for hierarchical clustering.

### Referenced Datasets

We obtained microarray gene expression profiling data from the following GEO datasets (http://www.ncbi.nlm.nih.gov/geo/): GSE3790, which includes human post-mortem striatal tissue from 38 HD-gene-positive cases and 32 age- and sex-matched controls^14^ (more information in supplementary table 11). HD cases were analysed according to the presence or absence of disease symptoms and Vonsattel pathological grade (scale=0–4). According to the Vonsattel scale, two neuropathologists with special interest and expertise in HD pathology graded the corresponding formalin-fixed and paraffin-embedded caudate section of HD cases. This grading scale is based on the overall pattern of caudate nucleus neuropathology, the number and proportion of neurons and astrocytes. Some cases were cross-referenced to confirm the consistency of the scoring procedure. DNA was extracted from all patients and controls, and the CAG repeat length allele in IT15 (HD gene) was genotyped. The Affymetrix microarrays, we downloaded the gene expression matrix to identify the DEGs (adjusted P < 0.05) by using the limma package.

### Functional Annotation GO Term Analysis

Functional annotation for the expressed gene list was performed with the clusterProfiler, GO.db, DOSE, org.Mm.eg.db and org.Hs.eg.db packages in R. The adjusted p-value method was controlled by the BH algorithm. GO terms with adjusted p-values < 0.05 were judged to be significant terms. The enrichGO function was used to perform the GO enrichment and the enrichKEGG function was used to make KEGG enrichment.

### Two-Tailed Hypergeometric Statistical Test

Two-tailed hypergeometric statistical tests were carried out by R (the phyper function, lower.tail=F) to determine whether there was significant overlapping (p-value < 0.05) of the number of DEGs on a specific pathway between striatal development (P14 to P60) and HD. In the application of the phyper function, the total number of genes involved in the pathway was from the KEGG database.

### Weighted Gene Co-expression Network Analysis

Weighted gene co-expression network analysis was performed using the R package WGCNA ^15^. The log_2_ (FPKM value +1) transformed data were used to generate a matrix of the Pearson correlations between all pairs of genes across the samples. The value β = 13 was chosen as the saturation level for a soft threshold of the correlation matrix based on the criterion of approximate scale-free topology. To minimize the effects of noise and spurious connections, the correlation matrix was transformed to the topological overlap matrix using the “TOMsimilarity” function ^15^. The topological overlap matrix was then used to group highly co-expressed genes by performing average linkage hierarchical clustering. The dynamic hybrid tree-cut algorithm was used to cut the hierarchal clustering tree ^15^, and the modules were defined as the branches resulting from this tree cutting. The minimum module size was defined as 30, and the threshold for the merging of similar modules was defined as 0.35.

Then, to show trends in gene expression, the genes from each module were grouped into clusters using the Mfuzz package in R with the fuzzy c-means algorithm ^16^. Before clustering, standardization of the expression values of every gene was carried out, so that the average expression value for each gene was 0, and the standardization deviation of its expression profile was 1.

### Processing of Machine Learning

Genes that belong to the similarity functional network may be co-expressed in the temporal logic and, thus, should have similar expression profiles to a certain extent. The exploration of the similarity between gene expression profiles can be transformed into the exploration of the correlations between them. At the same time, the correlations between known functional gene sets can be taken as an internal reference. Through machine learning, it is possible to explore the similar functional networks and the diversity of gene functions to a certain extent. For now, machine learning of correlations between substances has been used to search for potentially relevant substances, such as the components of protein complexes in their natural state ^35,40,41^. Therefore, we hope to explore similar functional gene networks and gene functional diversity on a large scale to some extent.

We applied several methods to measure the similarity of two gene expression profiles to emphasize multiple gene expression profile features. We treated each expression profile as a vector consisting of the observed FPKM for a particular gene across the corresponding temporal logical, and a complete co-expression experiment was stored as a matrix in which the rows and columns represent genes and times, respectively. To measure the co-expression profile similarity between two genes, we employed various correlation metrics that ranged from simple scores to more sophisticated metrics based on information theory. The Jaccard score (Jaccard) computes the ratio of how often genes are expressed at the same time point and how often genes are detected at all times. Thus, the Jaccard score between two genes was calculated by counting the number of time points that contained both genes and dividing the result by the number of time points that included at least one of the two genes. The Pearson correlation coefficient (PCC) is used to measure the similarity of two gene co-expression profiles. The PCC was calculated using the vector of the raw FPKM. Mutual information (MI) considers both linear and nonlinear dependencies between vectors. The Bayes correlation (Bayes) was proposed to process RNA-seq gene expression data based on sequence counts for various genes under different conditions. Here, we applied the same method for the FPKM for various genes across the temporal logic. For the Apex score (Apex), most genes tend to be expressed at a specific time, and thus, the time point that contains the largest amount of a particular gene is typically also the most critical time for that gene. Thus, two genes are considered to be more likely to have similar functions if the time points containing the largest recorded amounts of these genes across all times are the same.

These correlation coefficients were all between 0-1 (except the Apex, which was either 0 or 1). To reduce the operation time, we reserved the pairs with the maximum correlation scores (gene co-expression pairs in all four different correlation measures) ≥ 0.75/0.85. Afterward, the correlation vectors were input into a supervised machine learning model (random forest algorithm) that was both trained to predict new gene pairs and benchmarked against reference positive (annotated) gene groupings (that is, the enrichment of each module of genes in the GO database) (Table S10) and negative gene groupings (that is, combinations of genes with distinct enrichments in each gene module). The additional supporting evidence (for example, functional interactions inferred from co-expression) was integrated from public sources TRRUST2 ^36^, thereby producing richer and more accurate interaction networks. Finally, network partitioning to define gene network membership was applied by ClusterONE ^20^. Afterward, to prove that the co-expressed gene networks obtained through machine learning had more biological significance, the functional significance of the randomly generated gene networks was compared. A randomly generated gene network is a random sampling of the genes involved in machine learning, and random sampling ensures that the number of gene networks and the total number of genes in each gene network are consistent. We conducted a total of 100 random samplings, conducted enrichment analysis of the results of each random sampling, and calculated its average frequency in each significance stage to exclude the contingency of random sampling.

### Static Network Analysis and Gene Regulatory Network Prediction

The reconstructed network was imported into the Cytoscape ^42^ software package for subsequent analyses. The network was analysed based on the topological parameters, i.e., the number of nodes, diameter, radius, centralization, density, heterogeneity, number of connected components, number of the shortest paths, characteristic path length, and average number of neighbours, and central parameters (i.e., the node degree distribution and neighbourhood connectivity distribution) using the Network Analyzer plug-in.

To infer the gene regulatory network, given the expression profiles and the list of genes, putative regulatory links from genes to all other genes were estimated using Genie3 ^43^ with default parameters, and only the top 25% of regulatory links with a prediction weight were retained for constructing a GRN. For each node (gene) in the network, the in- and out-degrees represented the number of edges (regulatory links) directed to and from this node, respectively.

### Annotation of Genes Using Existing Datasets

The transcription factor annotation integrated three databases (Table S10), which were TRRUST2, TFdb and GO. TRRUST2 (https://www.grnpedia.org/trrust/) is a manually curated database of human and mouse transcriptional regulatory networks^36^. TFdb (http://genome.gsc.riken.jp/TFdb/) is a database containing mouse transcription factor genes and their related genes. The genes are annotated in the Gene Ontology database as GO:0003700, “DNA binding TF activity”.

The disease annotation was from the DisGeNET (https://www.disgenet.org/) database, which is a discovery platform containing one of the largest publicly available collections of genes and variants associated with human diseases ^19^. The genes annotated in the DisGeNET database as Nervous System Diseases, Mental Disorders, Behavior and Behavior Mechanism were collected as custom nervous system-associated disease databases (Table S10).

The homologous annotation was from the Brain Atlas (https://www.proteinatlas.org/humanproteome/brain), which explores protein expression in the mammalian brain by the visualization and integration of data from three mammalian species (human, pig and mouse) ^18^.

## Acknowledgements

We are grateful to Guillermo Oliver at Northwestern University for the generous gift of Six3^F/F^ mice, and Kenneth Campbell at University of Cincinnati College of Medicine for the Dlx5/6-Cre mice.

## Data availability statement

The data that support the findings of this study are available from the corresponding author upon reasonable request.

